# Optimal drug treatment for reducing long-term drug resistance

**DOI:** 10.1101/2022.07.29.502041

**Authors:** Tina Ghodsi Asnaashari, Young Hwan Chang

## Abstract

The maximum-tolerated dose principle, the highest possible drug dose in the shortest possible time period, has been the standard care for cancer treatment. Although it is appealing in a homogeneous tumor settings, tumor heterogeneity and adaptation play a significant role in driving treatment failure. They are still major obstacles in cancer treatments despite great advances in modeling and cancer therapy using optimal control theory. To address this, we first generalize two population models and examine the long-term effects of differential selective treatment strategies. Second, we take into account different drug-imposed selective pressure into designing optimal treatment strategies. Numerical examples demonstrate that the proposed treatment strategy decreases long-term tumor burden by decreasing the rate of tumor adaptation.

## I. INTRODUCTION

In general, the application of optimal control theory in cancer treatment is proposing a mathematical model and optimization to find the best treatment method for preventing tumor growth by satisfying some desirable conditions [1]– [6]. Despite all these efforts to find the optimal regime of cancer treatment, drug treatment may lead to new mutations through both genetic and non-genetic mechanisms as the tumor gains drug resistance and this drug resistance is an impediment to treating cancer. Therefore, tumor heterogeneity and adaptation continue to be a major barrier to the successful treatment of cancer [7].

The maximum-tolerated dose (MTD) principle has been the standard care for cancer treatment for several decades and is the basis for clinical evaluation [8]. Although MTD therapy is appealing in a homogeneous tumor setting, it may not be the optimal control strategy for heterogeneous tumors or competition between sensitive and resistant cells. In [9], the authors also reported that MTD is not a good treatment strategy as it may lead to an increased resistant cell population. Since recent biological studies demonstrated the evidence of tumor heterogeneity and assessed potential biological and clinical implications [7], [10], [11], finding new strategies is still an open area of research to overcome tumor heterogeneity and adaptation.

To address tumor heterogeneity, recent studies proposed mathematical models to consider different cell population dynamics [8], [9], [12]–[15]. The simplest form of these models is those which use sensitive and resistant cells as two cell type species in an ordinary differential equation (ODE) model [16] and optimal control is designed to avoid the resistant population becoming dominant. In [8], evolutionary strategies were proposed with conventional therapies in order to control the appearance of drug resistance by considering spatial heterogeneity. In [17] and [18], the authors modeled tumor heterogeneity, with the presence of different cancer cell subpopulations each with a different level of susceptibility to each of the available drugs (and with the possible presence of metastasis in various body compartments where the drugs can have different efficacy), and then proposed optimal treatments in order to reduce the tumor size as well as possible side effects, also considering immunotherapy [19].

In [13], the authors modeled the long-term effects of two different drug treatment methods, symmetric and asymmetric treatment regimes in which the drug’s effect on the subpopulation is equal and different respectively. Selective treatment pressure is the influence exerted by drugs to promote one group of sub-population over another that may shift tumor heterogeneity distribution and generate resistance cells to the drug. The authors performed simulation studies with sensitivity analysis by using parameter sweeps to analyze the effects of each parameter on therapeutic efficacy and interrogate the effects of different drug-imposed selective pressures on long-term therapeutic outcomes. However, it is limited to drawing a fundamental and principled understanding of the effect of differential selective pressure.

In this paper, motivated by [13], we first generalize two population models and understand the effect of differential selective treatments. We examine different drug-imposed selective pressure effects and take into account this principle on optimal control design. Second, we formulate the optimal control problem to penalize a rate of tumor adaptation while minimizing tumor burden. We consider two different scenarios: 1) a single drug profile that has effects on the entire cell population at the same time and 2) multiple drug profiles where each drug only affects corresponding subpopulation on its drug cycle. We also consider different drug duration times for each drug profile in order to consider a more practical scenario. Finally, numerical simulations are introduced to demonstrate the proposed approach.

## II. Background

In this section, we briefly summarize the previous work [13] as we extend this study by focusing more theoretical analyses and providing a generalized model (i.e., considering more than two populations).

In [13], a minimal two-population was modeled as (*x*_1_, *x*_2_) with distinctive growth rates (*k*_1_,*k*_2_) and drug killing rates (*α*_1_,*α*_2_) respectively. The kinetics of the two subpopulations were modeled using a simple ODE for expo-nential growth as follows:

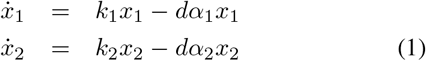

where drug treatment (*d*) is a Heaviside step function as shown in Figure 1. In the problem setting [13], the authors assumed the same initial overall tumor growth and tumor reduction for the first treatment (i.e., from 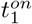 and 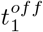 where 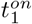 and 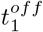 represent the start time point and the end time point of the first treatment respectively) of both symmetric and asymmetric treatment conditions in order to examine long-term effects of two different treatment regimes.

**Fig. 1.**
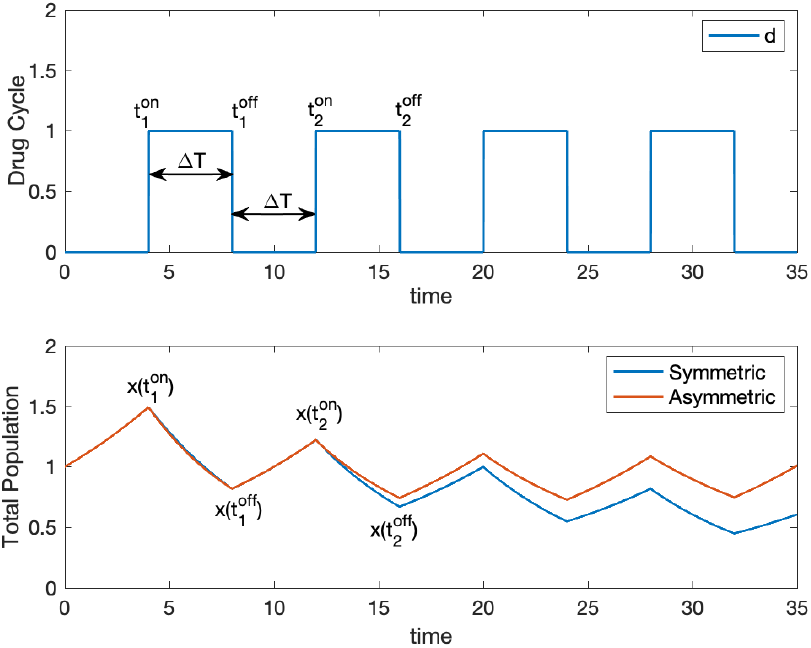
A comparison of total tumor population (i.e., *x* = *x*_1_ + *x*_2_) between symmetric and asymmetric treatment schemes; Although the overall population is the same at 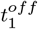 (the same initial and overall tumor size at the time of treatment and the same initial efficacy on the overall tumor), asymmetric treatment results in higher tumor burden in the long term compared to symmetric treatment. The top figure shows drug treatment cycle and the bottom figure shows the overall tumor population dynamics.

Thus, during the initial untreated growth phase of the tumor and after the first treatment, the total tumor size is equivalent to each other, for instance, the same killing effect on the different tumor cell types (symmetric) and different killing effects on the different tumor cell types (asymmetric) cases as shown in Figure 1. A simulation result showed that symmetric treatment is more effective than asymmetric treatment. This result shows that tumor burden decreases more when symmetric treatment is applied to the system.

## III. Differential-imposed selective treatments with a single drug profile

Motivated by the effects of distinct drug selective pressures on long-term tumor response [13], we generalize a two-population model and consider how to use this principled concept in treatment design that ultimately minimizes tumor burden. We consider a general case where we have *m* states as follows:

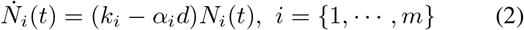

where *N_i_* represents the population of the i-th cell type and drug treatment (d) is a Heaviside step function where we simply assume 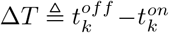 is equal to drug-off duration (i.e., 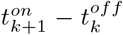) and constant over *k*. Then, we define a population composition rate:

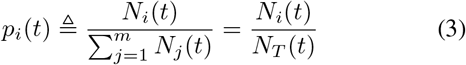

where 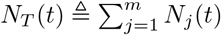 is the total population. The rate of composition change is as follows:

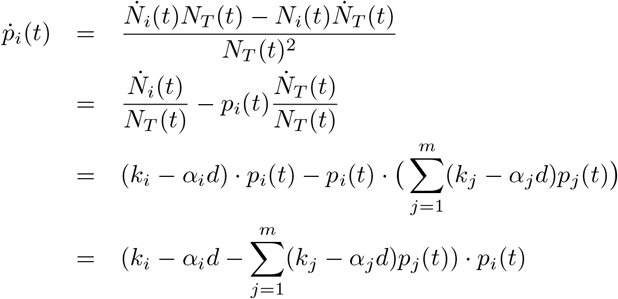

### Lemma 1

For symmetric treatment, sub-population composition does not change over time.

*Proof:* For symmetric treatment, we have *k_i_* — *α_i_ d* = *k_j_* — *a_j_d* where *i* ≠ *j*. Thus, we have the following:

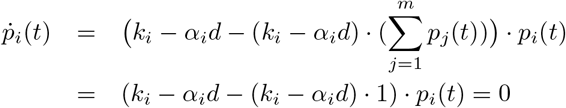

In Lemma 1, we showed that symmetric treatment condition guarantees 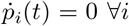 (i.e., sufficient condition). To show that it is a necessary condition for 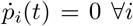, we prove the following lemma.

### Lemma 2

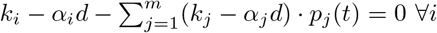 (i.e., 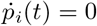), then *k_i_* – *α_i_d* = *k_j_* — *a_j_d* where *i* ≠ *j*.

*Proof:* (by induction)

Assuming that it is true for *m*, i.e., ∀*i* = {1, … *m*}, 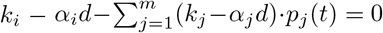 implies *k_i_* – *α_i_ d* = *k_j_* – *α_j_d* where *i* ≠ *j*. Then, we prove that it is true for *m* + 1:

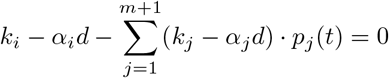

Rearranging this equation:

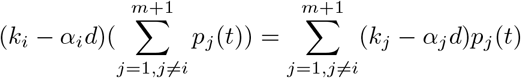

Using the assumption that 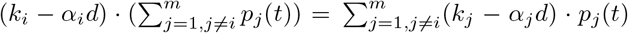 implies *k_i_* — *α_i_d* = *k_j_* – *α_j_d* where *i* ≠ *j* and *i* = {1, …, *m*}. Then, we have

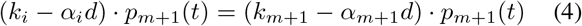

where *i* ≠ *m* +1 and thus *k_i_* – *α_i_d* = *k*_*m*+1_ – *α*_*m*+1_*d*

To measure long-term effects of treatment regimes, we define a tumor reduction rate as follows.

### Definition 1

A Tumor Reduction (TR) rate after each round can be defined as follows [13]:

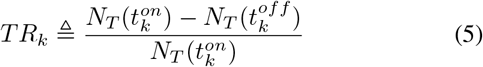

where *TR_k_* represents a tumor reduction rate of the *k*-th drug cycle, 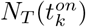 and 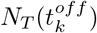 represent total tumor population at time step 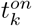 and 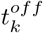 respectively as shown in Figure 1.

### Lemma 3

For symmetric treatment, a tumor reduction rate after each round is constant over time, i.e., *TR_k_* = *TR*_*k*+1_.

*Proof:* For a symmetric treatment, we have *k_i_* – *α_i_* = *k_j_* – *α_j_* = *k_s_* – *α_s_* and 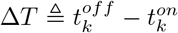 is assumed to be constant over *k*.

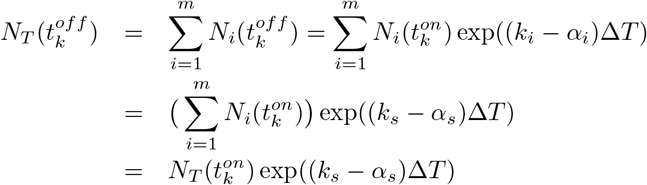

Therefore, for symmetric treatment, a tumor reduction rate is constant as follows:

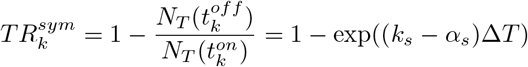

Similarly, we can show 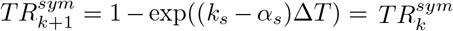.

### Lemma 4

For asymmetric case, a tumor reduction rate decreases over time, i.e., *TR*_*k*+1_ – *T*R_*k*_ < 0.

*Proof:* (suppose not) assume *TR*_*k*+1_ – *TR*_*k*_ ≥ 0:

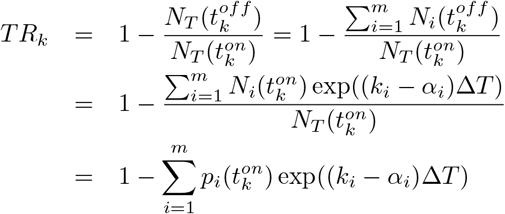

where we use 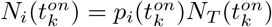.

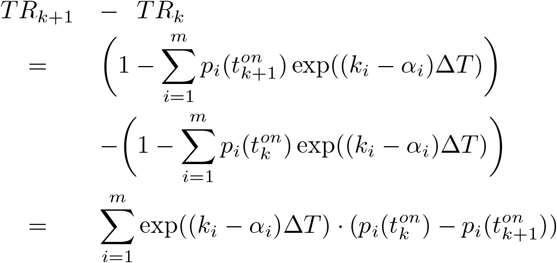

Without loss of generality, consider (*m* = 2):

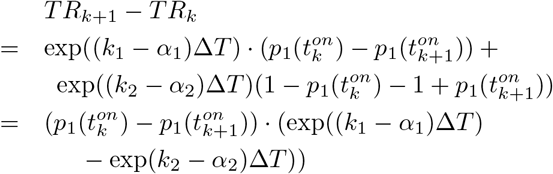

For asymmetric case (i.e., *k*_1_ – *α*_1_ ≠ *k*_2_ – *α*_2_), if *k*_1_ – *α*_1_ > *k*_2_ – *α*_2_, *p*_1_(*t*) increases as *p*_2_(*t*) decreases, i.e. 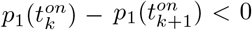 and thus *TR*_*k*+1_ – *TR*_*k*_ < 0 (contradiction). Similarly, if *k*_1_ – *α*_1_ < *k*_2_ – *α*_2_, *p*_1_(*t*) decreases as *p*_2_(*t*) increases, i.e. 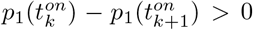 and then *TR*_*k*+1_ – *TR*_*k*_ < 0 (contradiction).

We also define the rate of tumor adaptation (or the rate of change in tumor sensitivity) by taking the slope of the percent tumor reduction values for successive doses.

### Definition 2

A rate of tumor adaption (TA) is defined as follows [13]:

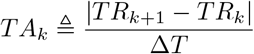

where *TR_k_* and *TR*_*k*+1_ represent tumor reduction at the *k^th^* and (*k* + 1)^*th*^ round of treatment respectively.

A rate of tumor adaptation refers to how quickly the tumor reduction changes by taking the difference between the absolute value of successive tumor reductions over time [13]. If the tumor adaptation rate increases, the effectiveness of the drug decreases by definition.

### Lemma 5

For symmetric treatment (i.e., equal selective pressure on treatment), a rate of tumor adaptation is zero (i.e., *TA_k_* = 0, ∀*k*).

*Proof:* By Lemma 3 and Definition 2.

### Theorem 1

With the same initial overall tumor size at the time of treatment and the same initial efficacy on the overall tumor, differential-imposed selective pressures on the individual sub-populations (i.e., asymmetric treatment) result in a higher tumor burden in the long term compared to symmetric treatment.

*Proof*: 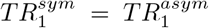 by assumption (i.e., the same initial efficacy on the overall tumor) and Lemma 4 and 5 (i.e., a tumor reduction rate is constant in symmetric treatment but decreases over time in asymmetric treatment).

Therefore, symmetric treatment minimizes the rate of tumor adaptation and thus maximizes the effectiveness of the drug over time.

As we cannot have control over subpopulation during the drug-off time period, now we consider drug on- and off-period as one treatment cycle (i.e., 2ΔT) and simply extend the condition to conserve sub-population in each drug cycle as follows:

### Lemma 6

In order to conserve subpopulation composition after one treatment cycle, i.e., 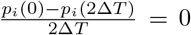, we need to satisfy 2*k_i_* – *α_i_* = 2*k_j_* – *α_j_* where *i* ≠ *j*.

*Proof:*

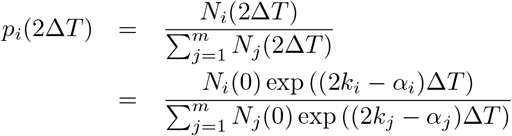

Thus, if 2*k_i_* – *α_i_* = 2*k_j_* – *α_j_* where *i* = *j*,

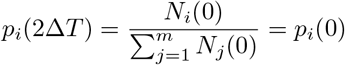

By considering one treatment cycle (i.e., drug-on and off), we could conserve sub-population especially when we have different growth rates for each cell population (i.e., *k_i_* ≠ *k_j_*). For example, when the drug is off (i.e., *d* = 0), different growth rates will change the composition rate as we do not have control but by considering the drug on/off cycle, we could compensate for composition change during the drug-off period and thus conserve subpopulation composition.

Now we define the objective function in the following form:

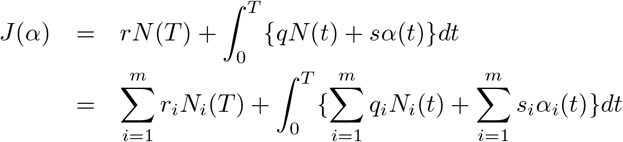

where *r, q* and *s* are constant parameters. Then the optimization problem can be described with these constraints to avoid increasing the rate of tumor adaptation and thus ultimately minimize tumor burden in the long term:

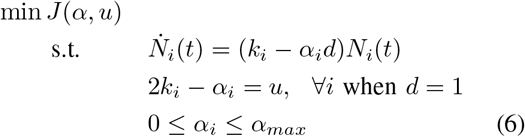

where we introduce another variable *u* as optimization variable to satisfy ∀*i*, 2*k_i_* – *α_i_* = 2*k_j_* – *α_j_* (*i* ≠ *j*). Note that we also consider the maximum drug effect (*α_max_*) as inequality conditions. By solving the optimization problem (6), we minimize the overall tumor burden while maintaining sub-population composition in order to minimize tumor adaptation.

## IV. Differential-imposed selective treatments with multiple drug profiles

In the previous section, we simply consider a single drug profile that can affect all sub-population together with different drug killing rates (*α_i_*). However, this is not feasible in a practical scenario and thus we consider multiple drug profiles where each drug profile has a single effect on the specific sub-population as shown in Figure 2. Similar to the previous section, we consider one full drug treatment cycle to conserve sub-population proportion.

**Fig. 2.**
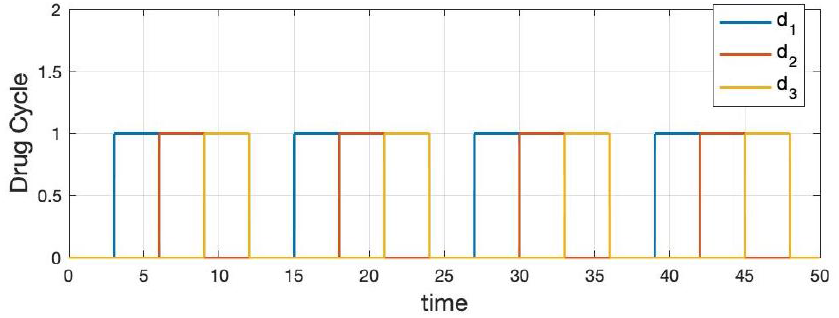
Treatments with multiple drug profiles (*m* = 3) where each drug has only effect on the corresponding sub-population.

### Lemma 7

Consider *n* drug profiles where the time interval for each drug is equal to the drug-off time interval (Δ*T*).

Sub-population proportion is conserved after one drug cycle if (*n* + 1) · *k_i_* – *α_i_* = (*n* + 1) · *k_j_* – *a_j_, i* ≠ *j*

*Proof:* Each sub-population is under the effect of the corresponding drug during Δ*T* and it increases with the growth rate (*k_i_*) when the drug is an off cycle (i.e., *n* · Δ*T*). So, after the whole drug cycle ((*n* + 1) · Δ*T*), the population of the *i*-th cell type can be represented by *N_i_*((*n* + 1) · Δ*T*) = *N_i_*(0) exp(((*n* + 1) · *k_i_* – *α_i_*) · Δ*T*). Thus,

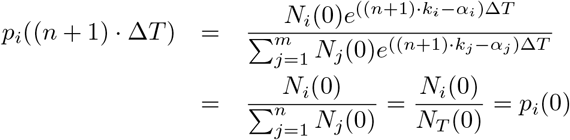

Therefore, after one full treatment cycle, we can conserve subpopulation composition, i.e., *p_i_*((*n* + 1) · Δ*T*) = *p_i_* (0) with multiple drug profiles.

With multiple drug profiles, the optimization problem can be described with these constraints to avoid increasing the rate of tumor adaptation in the long term:

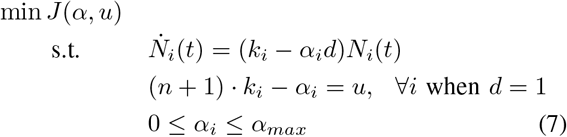

Thus, we minimize the overall tumor burden while maintaining sub-population composition in order to minimize tumor adaptation.

So far we simply consider that duration of each drug is constant (i.e., Δ*T*) but we also consider different duration times for each drug (i.e., Δ*T_j_* denotes the duration of the *j*-th drug and Δ*T*_0_ represents a duration of the drug-off period) with a simple extension as follows.

### Lemma 8

Consider n drug profiles with different drug duration (Δ*T* ≠ Δ*T_j_*). Sub-population proportion is conserved after one cycle if 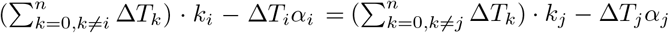.

*Proof:* Consider 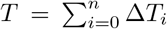 as duration of one drug cycle of multiple drugs treatment.

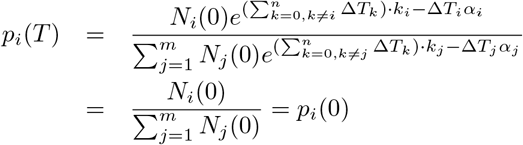

Finally, for multiple drug profiles and different treatment intervals, we consider the following optimization:

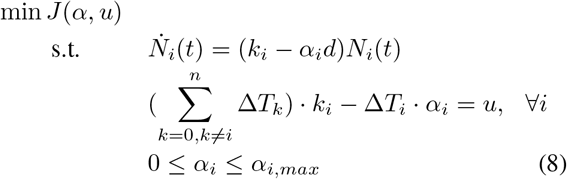

Note that we also consider different maximum drug effects for individual drug (*α_i, max_*).

## V. Results and Discussion

In this section, we demonstrate the effects of drug selective pressure with numerical simulations by solving the optimization problems (6) and (8). As we extended a minimal two-population case, we demonstrate three sub-population (i.e., *m* = 3) with a single drug profile and multiple drug profiles. For the first case, we simply consider the duration of each drug and drug-off duration equal to Δ*T* and for the second case, we consider different drug duration times. For each simulation study, we compare the overall tumor population and tumor reduction with and without conserving subpopulation (i.e., the proposed constraint in optimization).

### A. Example 1: with a single drug profile

For single drug profile study, we use parameters *k*_1_ = 0.1,*k*_2_ = 0.12,*k*_3_ = 0.14,*x*_1_ (0) = 0.45,*x*_2_ (0) = 0.30, *x*_3_(0) = 0.25, *α_max_* = 1 and *α_i_* where *i* = {1,2,3} are optimization variables. Figure 3 shows optimization result for single drug profile. After one drug cycle (i.e., drug-on and off denoted by a green dash line), the subpopulation ratio is conserved with the proposed constraint (blue line) and thus we observe that the proposed approach is more effective than the case without conserving subpopulation.

**Fig. 3.**
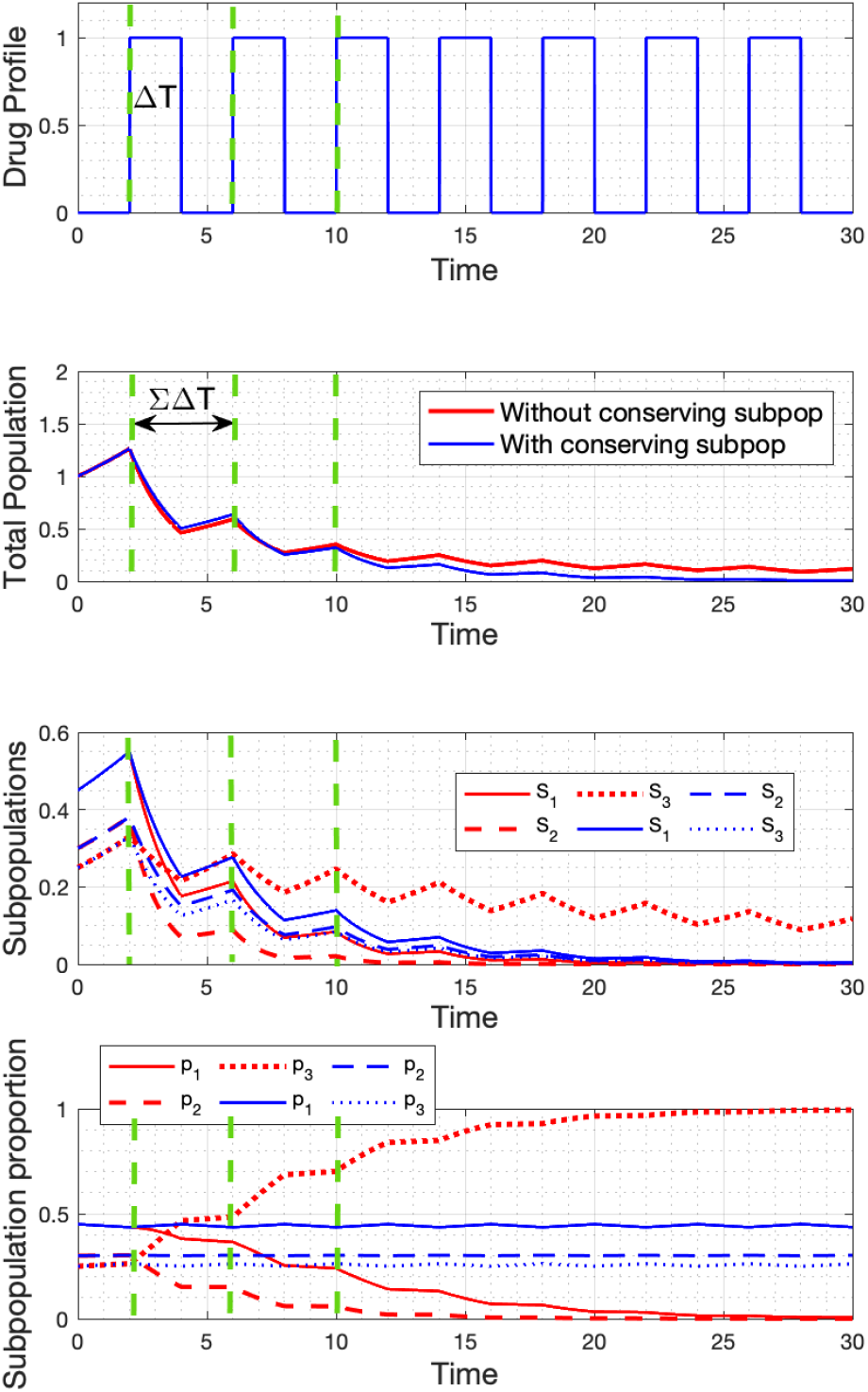
Simulation results with a single drug profile. By conserving subpopulation ratio, we maximize tumor reduction and thus have better longterm effects (blue). On the other hand, *p*_3_ population becomes dominant and shows resistant to therapy (red).

We observe that although some subpopulation decreases faster in the case without conserving subpopulation (red line), other sub-population (i.e., *p*_3_) remains as a resistant population to the drug and reduce long-term drug effects. Thus, conserving subpopulation-imposed treatment shows better long-term effects compared with differential selective pressure treatment (i.e., without conserving subpopulation).

As we derived tumor reduction for symmetric and asymmetric treatment in Lemma 3 and 4, the proposed approach (i.e., conserving subpopulation ratio) guarantees that a rate of tumor adaptation is zero so that ultimately minimizes tumor burden in the long term as shown in Figure 4 (right). On the other hand, without the proposed approach, we observe that tumor reduction decreases over time due to tumor adaptation as shown in Figure 4 (left). Note that tumor reduction of the first drug cycle without conserving subpopulation ratio is higher than tumor reduction of the first drug cycle with the proposed approach but it decreases over time while tumor reduction is conserved with the proposed method.

**Fig. 4.**
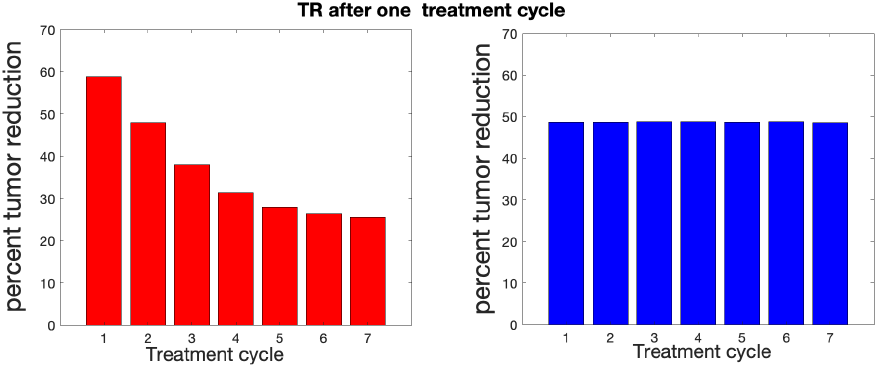
Tumor reduction after each round of drug treatment with a single drug profile. (left) Tumor reduction decreases when we have differential selective pressure on treatment. (right) On the other hand, the proposed approach guarantees that a rate of tumor adaptation is zero (i.e., tumor reduction is constant)

### B. Example 2: with multiple drug profiles and different drug duration time

We considered different drug duration for each drug (Δ*T_i_*) and drug-off duration (Δ*T*_0_) where we use Δ*T*, Δ*T*_2_, Δ*T*_3_ are 0.5, 1 and 2 respectively, Δ*T*_0_ = 2.5 and *α_i, max_* = 1. We consider optimization problem (8) and simulation result is shown as Figure 5. Again the proposed optimization can handle different drug duration times while conserving the subpopulation ratio. Thus, similar to the previous cases, the tumor reduction rate is conserved with the proposed approach, and tumor adaptation is minimized.

**Fig. 5.**
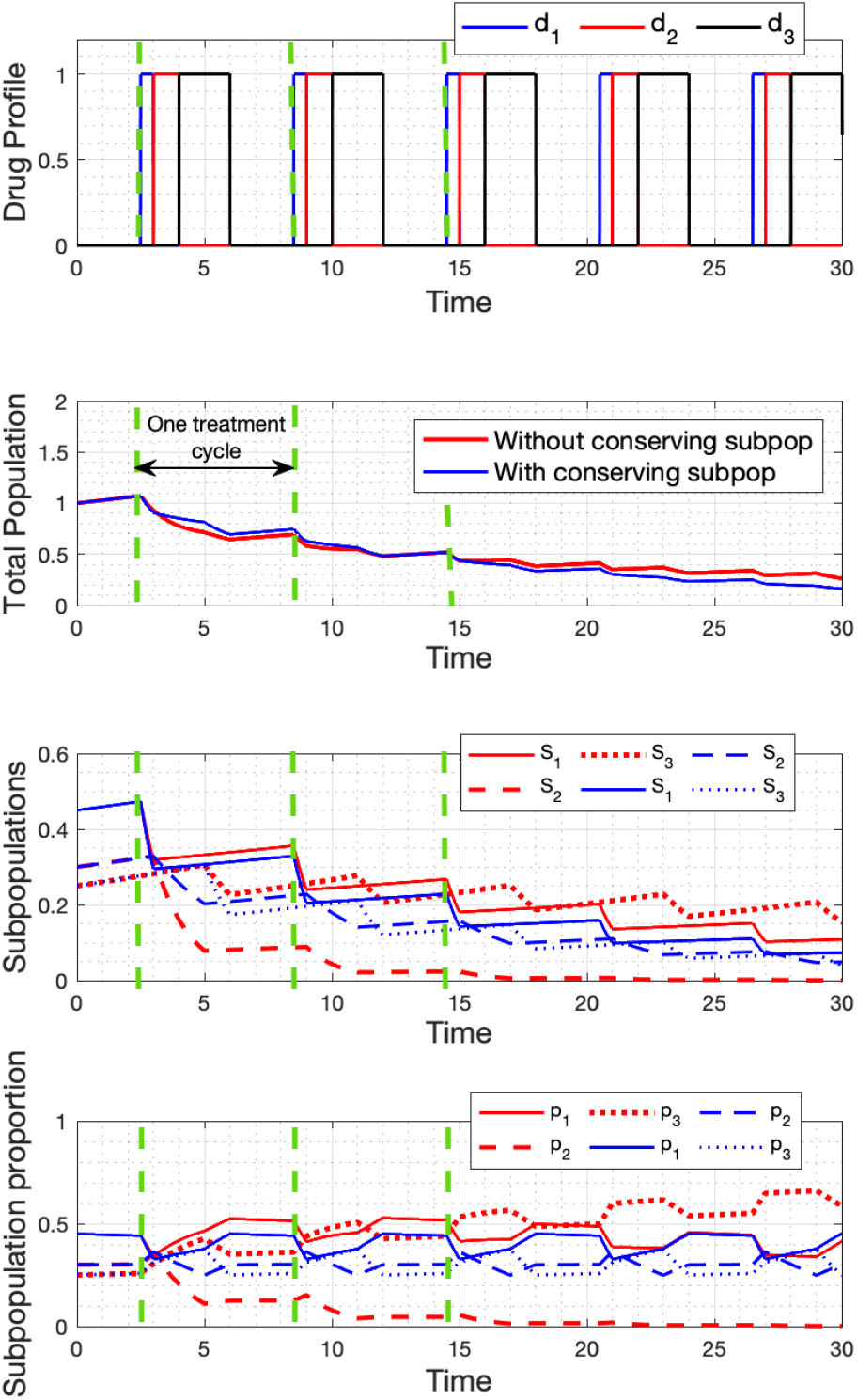
Simulation results with multiple drug profiles and different drug duration time (Δ*T_j_*).

## VI. Conclusion

In this paper, we consider tumor heterogeneity and selective pressure on sub-populations in the treatment design. Our simulations demonstrate the necessity of matching sub-population composition in order to minimize tumor adaptation. Our results indicate that different selective pressure on subpopulations causes the rate of composition change which can lead to the long-term effect of drug resistance and tumor adaptation. By conserving sub-population composition, we can minimize the rate of tumor adaptation and thus maximize a tumor reduction rate. We also show that sub-population composition change can cause decreasing a tumor reduction rate.

The tumor model presented here is greatly simplified but we have chosen a simple starting point to understand how different selective pressures affect tumor adaptation and subpopulation composition changes. Thus the model needs to be improved and modified to represent specific cancer types, resistance mechanisms, and tumor microenvironment factors in future work.

## ACKNOWLEDGMENTS

This work was supported in part by the National Cancer Institute (U54CA209988).

## References

[1] A. S. Matveev and A. V. Savkin, “Application of optimal control theory to analysis of cancer chemotherapy regimens,” Systems & control letters, vol. 46, no. 5, pp. 311–321, 2002.

[2] M. Leszczyński, U. Ledzewicz, and H. Schättler, “Optimal control for a mathematical model for chemotherapy with pharmacometrics,” Mathematical Modelling of Natural Phenomena, vol. 15, 2020.

[3] A. M. Jarrett, D. Faghihi, D. A. Hormuth, E. A. Lima, J. Virostko, G. Biros, D. Patt, and T. E. Yankeelov, “Optimal control theory for personalized therapeutic regimens in oncology: Background, history, challenges, and opportunities,” Journal of clinical medicine, vol. 9, no. 5, p. 1314, 2020.

[4] J. L. Boldrini and M. I. Costa, “Therapy burden, drug resistance, and optimal treatment regimen for cancer chemotherapy,” Mathematical Medicine and Biology: A Journal of the IMA, vol. 17, no. 1, pp. 33–51, 2000.

[5] L. G. de Pillis, W. Gu, K. R. Fister, T. Head, K. Maples, A. Murugan, T. Neal, and K. Yoshida, “Chemotherapy for tumors: An analysis of the dynamics and a study of quadratic and linear optimal controls,” Mathematical Biosciences, vol. 209, no. 1, pp. 292–315, 2007.

[6] S. I. Oke, M. B. Matadi, and S. S. Xulu, “Optimal control analysis of a mathematical model for breast cancer,” Mathematical and Computational Applications, vol. 23, no. 2, p. 21, 2018.

[7] N. El-Sayes, A. Vito, and K. Mossman, “Tumor heterogeneity: A great barrier in the age of cancer immunotherapy,” Cancers, vol. 13, no. 4, p. 806, 2021.

[8] J. A. Gallaher, P. M. Enriquez-Navas, K. A. Luddy, R. A. Gatenby, and A. R. Anderson, “Spatial heterogeneity and evolutionary dynamics modulate time to recurrence in continuous and adaptive cancer therapies,” Cancer research, vol. 78, no. 8, pp. 2127–2139, 2018.

[9] U. Ledzewicz, S. Wang, H. Schättler, N. André, M. A. Heng, and E. Pasquier, “On drug resistance and metronomic chemotherapy: A mathematical modeling and optimal control approach,” Mathematical Biosciences & Engineering, vol. 14, no. 1, p. 217, 2017.

[10] L. G. Martelotto, C. K. Ng, S. Piscuoglio, B. Weigelt, and J. S. Reis-Filho, “Breast cancer intra-tumor heterogeneity,” Breast Cancer Research, vol. 16, no. 3, pp. 1–11, 2014.

[11] A. Marusyk and K. Polyak, “Tumor heterogeneity: causes and consequences,” Biochimica et Biophysica Acta (BBA)-Reviews on Cancer, vol. 1805, no. 1, pp. 105–117, 2010.

[12] M. P. Chapman, T. Risom, A. J. Aswani, E. M. Langer, R. C. Sears, and C. J. Tomlin, “Modeling differentiation-state transitions linked to therapeutic escape in triple-negative breast cancer,” PLoS computational biology, vol. 15, no. 3, p. e1006840, 2019.

[13] D. Sun, S. Dalin, M. T. Hemann, D. A. Lauffenburger, and B. Zhao, “Differential selective pressure alters rate of drug resistance acquisition in heterogeneous tumor populations,” Scientific reports, vol. 6, no. 1, pp. 1–13, 2016.

[14] B. Zhao, M. T. Hemann, and D. A. Lauffenburger, “Intratumor heterogeneity alters most effective drugs in designed combinations,” Proceedings of the National Academy of Sciences, vol. 111, no. 29, pp. 10773–10778, 2014.

[15] B. Zhao, J. R. Pritchard, D. A. Lauffenburger, and M. T. Hemann, “Addressing genetic tumor heterogeneity through computationally predictive combination therapy,” Cancer discovery, vol. 4, no. 2, pp. 166–174, 2014.

[16] C. Carrère, “Optimization of an in vitro chemotherapy to avoid resistant tumours,” Journal of Theoretical Biology, vol. 413, pp. 24–33, 2017.

[17] G. Giordano, A. Rantzer, and V. D. Jonsson, “A convex optimization approach to cancer treatment to address tumor heterogeneity and imperfect drug penetration in physiological compartments,” in 2016 IEEE 55th Conference on Decision and Control (CDC), pp. 2494–2500, IEEE, 2016.

[18] C. A. Devia and G. Giordano, “Optimal duration and planning of switching treatments taking drug toxicity into account: a convex optimisation approach,” in 2019 IEEE 58th Conference on Decision and Control (CDC), pp. 5674–5679, IEEE, 2019.

[19] N. Dullerud and V. D. Jonsson, “Cellular immunotherapy treatment scheduling to address antigen escape,” in 2020 59th IEEE Conference on Decision and Control (CDC), pp. 4634–4639, IEEE, 2020.

